# CD81 represses NF-κB in HCV-expressing hepatoma cells

**DOI:** 10.1101/2023.09.13.557511

**Authors:** Maximilian Bunz, Mona Eisele, Dan Hu, Michael Ritter, Julia Kammerloher, Sandra Lampl, Michael Schindler

**Author notes:** corresponding author, michael.

## Abstract

The tetraspanin CD81 is one of the main entry receptors for Hepatitis C virus, which is a major causative agent to develop liver cirrhosis and hepatocellular carcinoma (HCC). Here, we identify CD81 as one of few surface proteins that are downregulated in HCV expressing hepatoma cells, discovering a functional role of CD81 beyond mediating HCV entry. CD81 was downregulated at the mRNA level in hepatoma cells that replicate HCV. Kinetics of HCV protein expression were increased in CD81-knockout cells and accompanied by enhanced cellular growth. Furthermore, loss of CD81 compensated for inhibition of pro-survival TBK1-signaling in HCV expressing cells. Analysis of functional phenotypes that could be associated with pro-survival signaling revealed that CD81 is a negative regulator of NF-κB. Interaction of the NF-κB subunits p50 and p65 was increased in cells lacking CD81. Similarly, we witnessed an overall increase in the total levels of phosphorylated and cellular p65 upon CD81-knockout. Finally, translocation of p65 in CD81-negative hepatoma cells was markedly induced upon stimulation with TNFα or PMA. Altogether, CD81 emerges as aregulator of pro-survival NF-κB signaling. Considering the important and established role of NF-κB for HCV replication and tumorigenesis, the downregulation of CD81 by HCV and the associated increase in NF-κB signaling might serve as viral mechanism to maintain persistent infection, ultimately causing chronic inflammation and HCC.

**Highlights:**

- CD81 is downregulated and transcriptionally silenced upon HCV genome replication
- Loss of CD81 is associated with increased cell growth and HCV expression
- CD81 suppresses NF-κB signaling.
- CD81 interferes with p65 activation and nuclear translocation

## Introduction

Liver-related diseases are responsible for approximately 2 million deaths annually (1–3). Of those, an estimated 300,000 were caused by hepatitis C virus (HCV) in 2019 (4). However, acute infection is not the major cause of HCV-related deaths, but liver cirrhosis and hepatocellular carcinoma (HCC) that can develop during chronic HCV infection (5). With no vaccine available and highly effective therapy options accessible to only a minority of the world’s population, greater efforts are required to decrease HCV-related disease burden (4,6,7).

Many aspects of HCV molecular biology have been elucidated within the last decades, but the changes in cellular homeostasis during chronic HCV infection are less well understood. It is well known that fibrogenesis and continuous inflammation of the liver are prerequisites for cancer development (5,8,9). Several studies have found that chronic HCV infection leads to chronic liver inflammation (5,8,9). However, the underlying mechanism that promotes the transition to cirrhosis and cancer has not been identified yet. One candidate described in the literature is cellular stress, which has been shown in some studies to be increased in patients with chronic HCV infection (10–12). Other studies found a general dysregulation of pathways that are associated with cancer development such as the cell cycle, DNA repair, pro-survival signaling and apoptosis (13–15).

The tetraspanin family of proteins is well known to serve as scaffolds for cell surface signaling complexes, for example, in the immunological synapse (16,17). Tetraspanins consist of four transmembrane helices connected by three domains: a large and a small extracellular loop (LEL/SEL), plus a short intracellular loop (16,17). The four transmembrane helices can form a cavity that binds cholesterol which can induce conformational changes (18). Several tetraspanins have been connected to viral-related processes such as entry (CD151 for HPV) or budding (CD63 and CD81 for HIV) (19). CD81 is also a cellular receptor for HCV that is bound by the viral E1/E2 glycoprotein complex and mediates entry (20,21). Furthermore, CD81 is involved in several signaling events, such as B cell receptor signaling through interaction with CD19, NK cell activation via with Adhesion G-protein coupled receptor G1, and presumably EGFR signaling (22–26).

In a screening approach to characterize cell surface receptor modulation in HCV-replicating hepatoma cells, we identified two tetraspanins, CD63 and CD81, as proteins that were downregulated. Based on this, we here characterized the mechanism of HCV-mediated tetraspanin modulation and analyzed the functional role of CD63 and CD81 in HCV-infected cells.

## Materials and Methods

### Cell culture

HEK293T and HeLa cells were cultured in DMEM (Thermo Fisher) supplemented with 10% fetal calf serum (FCS; Thermo Fisher) and 1% Penicillin/Streptomycin (Life Technologies). Huh7.5 and Huh7-Lunet cells, originally obtained from Charles Rice (Rockefeller University, New York), were cultured in DMEM (Thermo Fisher) supplemented with 5% fetal calf serum (FCS; Thermo Fisher), 1% Penicillin/Streptomycin, 1% Non-essential amino acids and 1% Sodium pyruvate (all Life Technologies). Stably transduced cells were cultured with additional 1 µg/ml puromycin. Cells were serum starved by culturing them in medium without FCS.

### LEGEND surface expression screen

To assess cell surface receptor modulation, the LEGENDScreen™ Human Cell PE Kit (Biolegend) was used. Huh7.5 cells were electroporated with Jc1_NS5A-mtagBFP. 48 h later, cells were detached and washed, before they were ntibody stained (5×10^4^-2×10^5^ cells per well). The staining and fixing procedure was performed as described previously (27).

### Plasmids and cloning

Plasmids were amplified in chemocompetent NEB10 or NEB Stbl3 (for CRISPR constructs) *E. coli* and isolated using the PureYield™Plasmid Midiprep System (Promega) according to the manual.

To generate pFK_Jc1_NS5A-mScarlet, the eGFP fluorescent protein of pFK_Jc1_NS5A-GFP (28) was replaced by mScarlet. In brief, the mScarlet insert was amplified from pmScarlet-C1 (Addgene #85042) with primers adding XbaI (XbaI-mScarlet_fw; 5’-GTtctagaCCTCGAGCT**ATGGTGAGCAAGGGCGA**-3’) and PmeI (meI-mScarlet_rev; 5’-CACgtttaaacCC**CTTGTACAGCTCGTCCATGC**-3’) restriction sites at the 5’- and 3’-ends, respectively. pFK_ Jc1_NS5A-GFP was digested with XbaI, PmeI and FastAP (Thermo Fisher) according to manufacturer’s instructions, separated by agarose gel electrophoresis, and the backbone band was cut out and isolated using NucleoSpin Gel and PCR cleanup Kit (Macherey-Nagel). Backbone and insert were ligated using T4 Ligase (Thermo Fisher) for 1 h at RT. Next, NEB10 chemocompetent bacteria were transformed with the ligation mix and plated on LB agar with 100 µg/ml ampicillin. Colonies were picked and a 5 ml culture was grown over night, followed by plasmid isolation (GeneJET Plasmid Miniprep System; Thermo Fisher). Isolated plasmids were test digested and sequenced.

pFK_Jc1_E2-mScarlet was generated according to Lee *et al.* (29), using a HCV genome with E2 N-terminally tagged with GFP. To generate a corresponding mScarlet expressing viral genome, a nucleotide sequence was synthesized (Genescript) starting at the Pfl23II restriction site in the E1 coding region, encoding mScarlet between E1 and E2 with the 3C peptide sequence connecting mScarlet and E2, flanked by XbaI. We additionally introduced an EcoRI restriction site between E1 and mScarlet and a BglII restriction site between mScarlet and the 3C peptide. The DNA construct the vector backbone (Jc1_E1(A4)_XbaI; similar to (30)) were cleaved with Pfl23II and XbaI (Thermo Fisher) and ligated as described above.

The fluorescent reporter construct where a fluorescent protein is N-terminally attached to the core coding region via a 2A self-cleaving peptide (pFK_Jc1_mScarlet-2A) analogous to the already described pFK_Jc1_R2A genome (31) was generated via restriction-free cloning. The principle is described at https://www.rf-cloning.org/ and primers were designed according to this protocol (32). In brief, primers were designed that where half complementary to the plasmid insertion site, and half complementary to mScarlet (Table 1). A PCR was performed with the insert pFK_Jc1_E2-mScarlet as template to generate a megaprimer such as that the mScarlet is flanked by EcoRI and BglII restriction sites. After purification of the megaprimer, it was mixed with the target plasmid, and a rolling circle PCR was performed to generate a plasmid with the insertion. The template was digested with DpnI (NEB) and the new viral genome was transformed into bacteria. To increase the chance of successful rolling circle PCR, a truncated version of pFK_Jc1_R2A was generated that only encoded the HCV genome until the end of E2. For this, pFK_Jc1_R2A was digested with SdaI and Pfl23II, and the insert was purified for ligation into pFK_Jc1_p7-half (30) to generate pFK_Jc1_R2A_p7-half. Then, pFK_Jc1_R2A_p7-half was digested with BcuI, HindIII and XbaI (Thermo Fisher), and the longest fragment (containing the backbone plus the HCV genome until the end of E2) was purified. Then BcuI and XbaI matching overhangs were ligated, giving rise to Jc1_R2A_short, which was then used as template for the rolling circle PCR. The HCV genome with the new reporter gene (pFK_Jc1_mScarlet-2A_short) was then digested with SdaI and Pfl23II and backligated into a full genome context using pFK_Jc1_R2A as backbone.

**Table 1:**
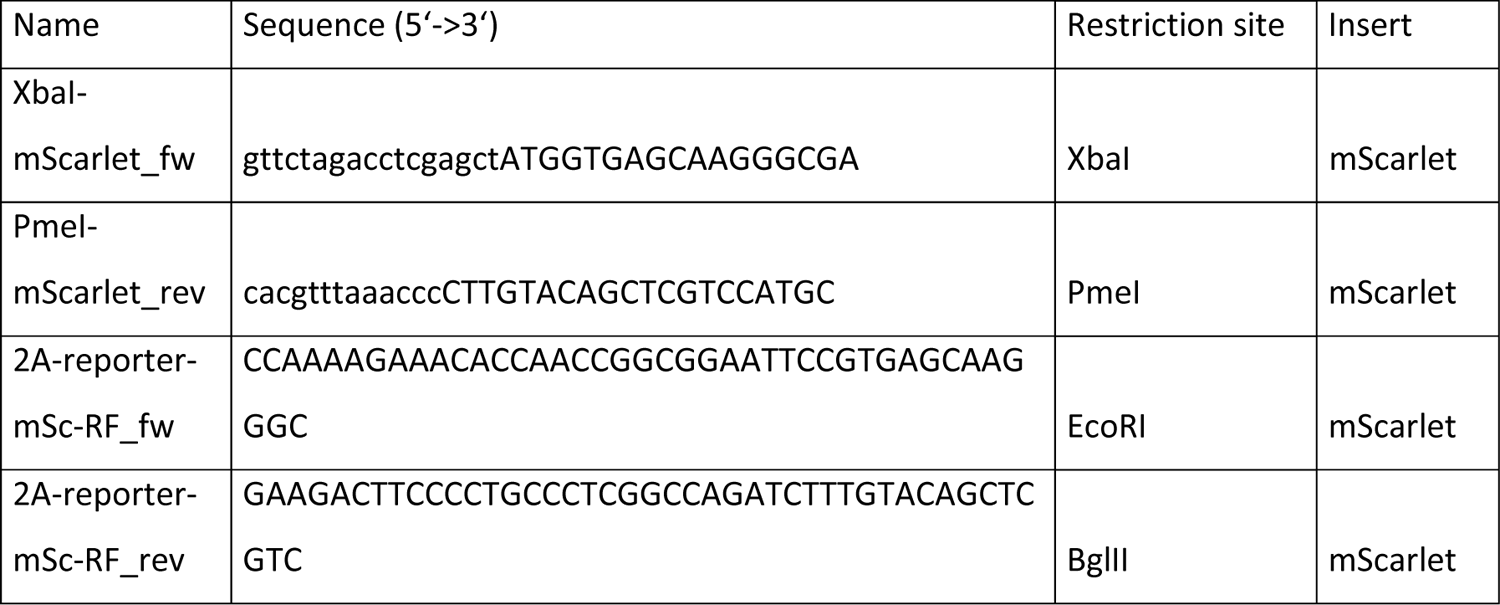
Cloning primers.

**Table 2:**
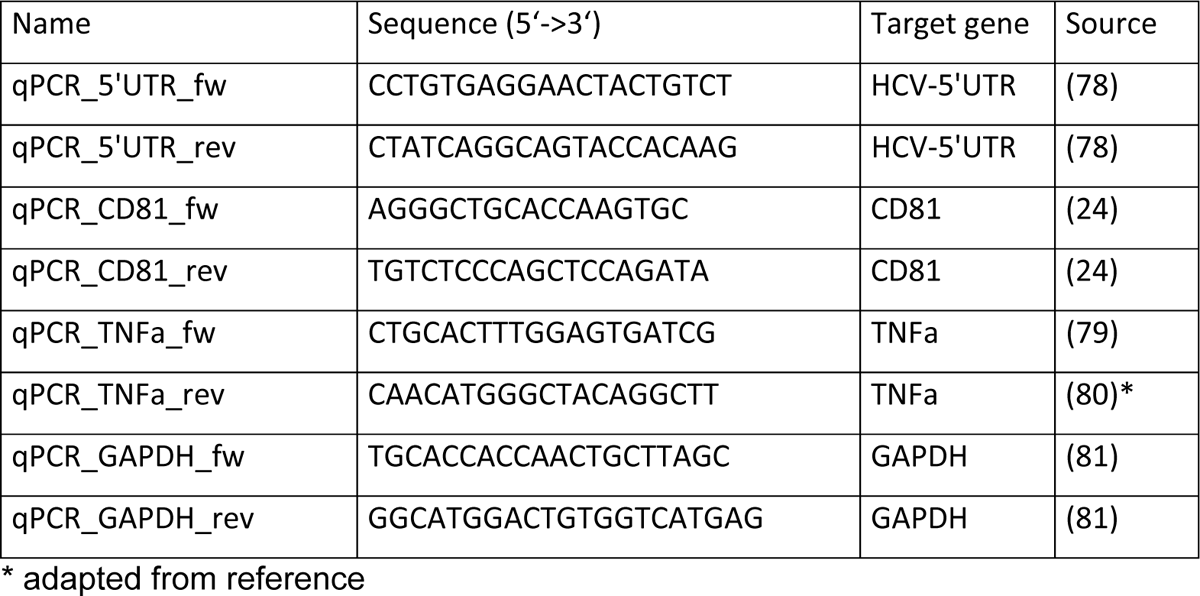
qRT-PCR primers.

**Table 3:**
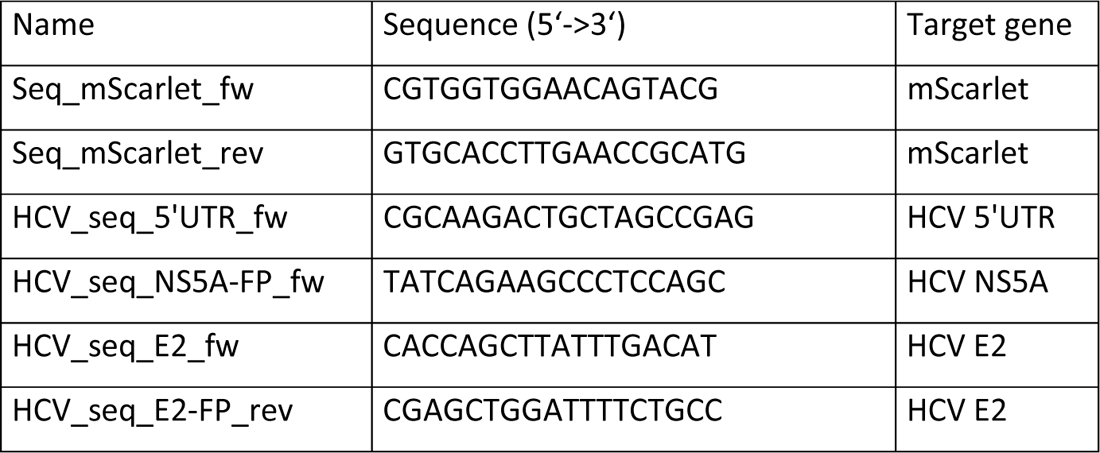
Sequencing primers.

### CRISPR/Cas9 plasmids and lentiviral production

To generate 293T, HeLa or Huh7.5 knock-out cells for tetraspanins CD63 and CD81, the LentiCRISPRv2 plasmid was used as described (33,34). In brief, oligonucleotides with the targeting sequence and specific overhangs for ligation were ordered (Metabion international); complementary oligonucleotides were annealed and phosphorylated, and then ligated into the LentiCRISPRv2 vector. The targeting sequences 5‘-GAGGTGGCCGCAGCCATTGC-3‘ (CD63) and 5’-CATCGGCATTGCTGCCATCG-3’ (CD81) were used. Subsequently, respective LentiCRISPRv2 constructs (3 µg/well) were transfected with lentiviral packaging (psPAX2; 2.25 µg/well) and envelope plasmids (pMD2G; 0.9 µg/well) in HEK293T cells using JetPRIME transfection reagent (Polyplus) according to the manufacturer’s instructions in a 6-well format. LentiCRISPRv2 without an integrated targeting sequence was used as control. 24-36 h after transfection, supernatant was harvested and spun at 3200 g for 10 min at RT to get rid of cellular debris. Cells were incubated with spun supernatant for 24 h and selected with puromycin (1 µg/ml) for 2 weeks.

### DNA transfection

HEK293T cells were transfected using polyethylenimine (PEI). Cells were seeded 24 h prior to transfection until they reached a confluency of 70-80%. In brief and exemplarily for a 12-well format, 1-2 µg plasmid DNA were added to 50 µl OptiMEM (Thermo Fisher) and another 50 µl PEI mix (OptiMEM with double the amount of PEI than DNA) was added. The transfection mixture was vortexed, spun down and incubated for 15 min at RT. 100 µl of the transfection mixture was dropped onto cells, and a medium change was performed 4-6 h later or the next day. DNA and PEI amounts were adjusted accordingly for transfection in other well formats. HeLa and Huh7.5 were transfected using JetPRIME transfection reagent (Polyplus) according to manufacturer’s instructions.

### *In vitro* viral RNA transcription

For *in vitro* transcription of viral RNA, the respective DNA vector was linearized by digestion using MluI (Thermo Fisher) for 1 h at 37°C (Table 4). Linearized vector was then purified using the Wizard® DNA Clean-Up System (Promega) according to manufacturer’s instructions. Complete linearization was checked by agarose electrophoresis, and DNA concentration was measured. 1 µg linearized vector was used for *in vitro* transcription using the T7 RiboMAX™Express Large-Scale RNA Production System (Promega) according to the manual. After DNA vector digest, a phenol chloroform RNA extraction was performed. Samples were filled up to 200 µl with nuclease-free water and 200 µl phenol:chloroform:isoamylalcohol (25:24:1; Thermo Fisher) was added, then vortexed for 1 min and spun at max speed for 2 min. The upper phase was transferred to a new tube, and 200 µl chloroform:isoamylalcohol (24:1; Sigma Aldrich) was added. Again, the sample was vortexed for 1 min, spun at max speed for 2 min, and the upper phase was transferred to a new tube. Subsequently, 20 µl 3 M sodium acetate (pH 5.2) and 200 µl isopropanol were added and the sample placed in ice for 5 min. Then, the sample was spun for 10 min at max speed to pellet the RNA, supernatant was discarded and the pellet washed with 70% EtOH. Finally, the pellet was dried at 37°C for 5 min and resuspended in 40 µl nuclease-free water. RNA was stored at −80°C.

**Table 4:** Viral genome constructs.

| Name | Restr. Sites used | Source |
| --- | --- | --- |
| pFK_Jc1_E1(A4) |  | (82) |
| pFK_Jc1_XbaI |  | (30) |
| pFK_Jc1_E1(A4)_XbaI |  | pFK_Jc1_E1(A4) |
| pFK_Jc1(A4)_mScarlet-HRV3C-E2 | SdaI, Pfl23II | pFK_Jc1_E1(A4),<br>based on (29) |
| pFK_Jc1_p7-half |  | (30) |
| pFK_Jc1_R2A_p7-half | SdaI, Pfl23II | pFK_Jc1_p7-half |
| pFK_Jc1_R2A |  | (31) |
| pFK_Jc1_R2A_short | BcuI, XbaI | pFK_Jc1_R2A_p7-half |
| pFK_Jc1_mScarlet-2A_short | RF cloning | pFK_Jc1_R2A_short |
| pFK_Jc1_mScarlet-2A | SdaI, Pfl23II | pFK_Jc1_R2A |
| pFK_Jc1_NS5A-GFP |  | (28) |
| pFK_Jc1_NS5A-mScarlet | PmeI, XbaI | pFK_Jc1_NS5A-GFP |

**Table 5:** Plasmids used for flow cytometry-based FRET.

| Name | Protein | Tag | Source |
| --- | --- | --- | --- |
| pECFP-C1 | eCFP |  | (77) |
| pECFP-C1 HCV Core | Core | eCFP | (77) |
| pECFP-N1 HCV E1 | E1 | eCFP | (77) |
| pECFP-C1 HCV E2 | E2 | eCFP | (77) |
| pECFP-C1 HCV p7 | p7 | eCFP | (77) |
| pECFP-C1 HCV NS2/3 | NS2-3 | eCFP | (77) |
| pECFP-N1 HCV NS3 | NS3 | eCFP | (77) |
| pECFP-C1 HCV NS4A | NS4A | eCFP | (77) |
| pECFP-C1 HCV NS4B | NS4B | eCFP | (77) |
| pECFP-N1 HCV NS5A | NS5A | eCFP | (77) |
| pECFP-C1 HCV NS5B | NS5B | eCFP | (77) |
| pEYFP-N1 | eYFP |  | (77) |
| pEYFP-N1-ECFP | eYFP-eCFP |  | (77) |
| pEYFP-C1 HCV Core | Core | eYFP | (77) |
| pEYFP-N1 HCV E1 | E1 | eYFP | (77) |
| pEYFP-C1 HCV E2 | E2 | eYFP | (77) |
| pEYFP-C1 HCV p7 | p7 | eYFP | (77) |
| pEYFP-C1 HCV NS2-3 | NS2-3 | eYFP | (77) |
| pEYFP-C1 HCV NS3 | NS3 | eYFP | (77) |
| pEYFP-C1 HCV NS4A | NS4A | eYFP | (77) |
| pEYFP-C1 HCV NS4B | NS4B | eYFP | (77) |
| pEYFP-C1 HCV NS5A | NS5A | eYFP | (77) |
| pEYFP-C1 HCV NS5B | NS5B | eYFP | (77) |
| pEYFP-C1 **CD63** | CD63 | eYFP |  |
| pEYFP-C1 **CD81** | CD81 | eYFP |  |

**Table 6:** Other plasmids.

| Name | Protein | Tag | Source |
| --- | --- | --- | --- |
| pWPI_BLR |  |  | (83) |
| pWPI-hCD81-HAHA-BLR | CD81 | HA-HA | (83) |
| pNFkB(3x)-Firefly Luciferase | FLuc |  | Daniel Sauter |
| pCMV-Gluc | GLuc |  | Daniel Sauter |
| pcDNA3.1 |  |  |  |
| p_human IKKβbeta, const. act. | IKKβ | | Daniel Sauter |
| p(N)FLAG-CMV2 MAVS | MAVS | Flag | Daniel Sauter |
| pEF-Bos-RIG-I 1-211-flag | RIG-I | Flag | Daniel Sauter |
| mKG_N |  | mKG_N |  |
| mKG_C |  | mKG_C |  |
| p50-mKG_N | p50 | mKG_N |  |
| p65-mKG_C | p65 | mKG_C |  |

### Electroporation of viral RNA

Electroporation was performed using the Neon Electroporation System (Thermo Fisher). For each electroporation 1-4×10^5^ (10 µl tip) or 1-4×10^6^ (100 µl tip) cells were used. Cells were seeded at the respective density 24 h prior to electroporation. Then, cells were detached and washed three times with PBS. Subsequently, cells were resuspended in an appropriate volume of PBS (containing Ca^2+^ and Mg^2+^; Thermo Fisher), and viral RNA was added (0.25-1 µg/1×10^6^ cells). The reaction chamber was filled with buffer E (buffer E2 for 100 µl tips). The cell/RNA mixture was put into an electroporation tip, and one pulse with 1300 V for 30 ms was applied. Electroporated cells were seeded accordingly in medium that did not contain any antibiotic.

### qRT-PCR

Cellular RNA of 2-5×10^5^ cells was extracted using the RNeasy Mini Kit (Qiagen) according to manufacturer’s instructions. For lysis, 1% 2-mercaptoethanol was added to the lysis buffer. 200 ng of extracted cellular RNA was used for cDNA transcription using the QuantiTect Reverse Transcription Kit (Qiagen) according to manufacturer’s instructions. cDNA samples were filled up to 60 µl with nuclease-free water. qRT-PCR measurements were carried out on a Lightcycler 480 (Roche) using Lightcycler 480 multiwell plates (Roche) and Luna Universial qPCR Master Mix (NEB) according to the manual. In brief, a master mix containing primers (final concentration 0.3 µM), Luna Universial qPCR Master Mix and nuclease free water was generated and added to the wells together with 2 µl of diluted cDNA in duplicates (Table 2). The ΔΔCp method was used for analysis.

### SDS-PAGE and western blot

Cells were lysed with RIPA buffer (10 mM Tris-HCl (ph 7.4), 1 mM EDTA, 0.5 mM EGTA, 140 mM NaCl, 0.1% (v/v) Na-deoxychalate, 0.1% (v/v) SDS, 1% (v/v) Triton X-100, 1x protease and phosphatase inhibitor) for 20 min at 4°C. Subsequently, 6x sample buffer (0.5 M Tris (pH 6.8), 0.6 M DTT, 30% (v/v) glycerol, 10% (w/v) SDS, 2% (w/v) bromphenol blue) was added accordingly, and the lysate was heated to 95°C for 10 min. Then, samples were loaded on a 12% acrylamide gel and separated at 80-140 V for 90-150 min. Blotting was performed in a wet blotting chamber (BioRad) at 80 V for 90 min. Then, the membrane was blocked for 1 h at RT using 5% milk or 1% BSA in PBS or TBS. Primary antibodies were applied over night at 4°C. Secondary antibodies were applied 1h at RT. Between steps, the membrane was washed three times with PBS-T or TBS-T (PBS or TBS with 0.1% Tween 20; Table 7). An Odyssey Fc Imaging System (LI-COR Biosciences) was used for visualization.

**Table 7:**
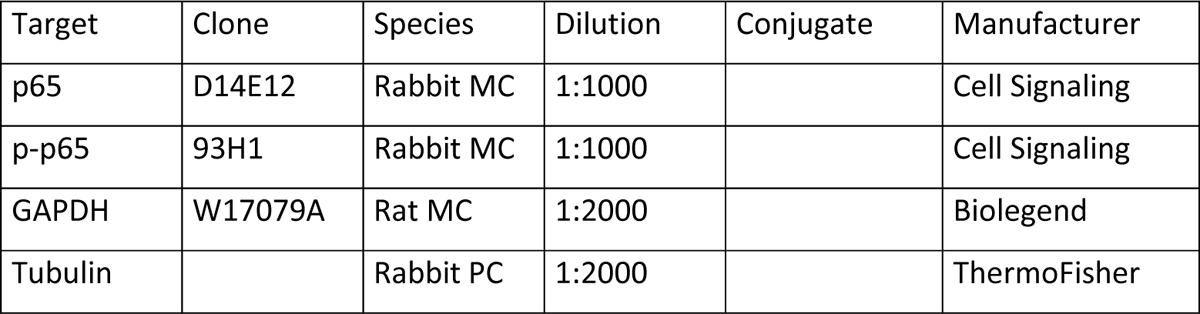
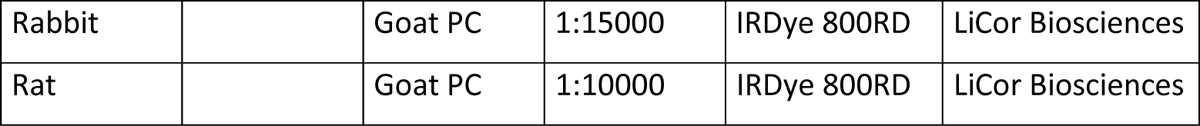
Anitbodies for western blot.

### Flow cytometry

For flow cytometry, cells were detached, washed and fixed with 2% PFA for 10 min at RT. For intracellular staining, cells were permeabilized with 0.2% saponin in PBS or 80% acetone for 10 min at RT. Staining with primary antibodies was performed for 30min at 4°C, followed by two washing steps with FACS buffer (1% FCS in PBS). If applicable, staining with secondary antibody was performed for 30 min at 4°C in the dark, also followed by two washing steps with FACS buffer (Table 8). Measurements were performed using a MACSquant VYB flow cytometer (Miltenyi Biotech).

**Table 8:** Antibodies for flow cytometry and immunofluorescence.

| Target | Clone | Species | Dilution | Conjugate | Manufacturer |
| --- | --- | --- | --- | --- | --- |
| CD63 | H5C6 | Mouse MC | 1:250 | PE | Biolegend |
| CD81 | 5A6 | Mouse MC | 1:250 | PE | Biolegend |
| CD81 | 5A6 | Mouse MC | 1:250 |  | Biolegend |
| CD317 | RS38E | Mouse MC | 1:250 | PE | Biolegend |
| CD317 | E-4 | Mouse MC | 1:250 | AF488 | SantaCruz Biotechn. |
| core | C7-50 | Mouse MC | 1:250 |  | Novus Biologicals |
| p65 | D14E12 | Rabbit MC | 1:250 |  | Cell Signaling |
| Mouse |  | Goat PC | 1:250 | AF594 | ThermoFisher |
| Rabbit |  | Donkey PC | 1:250 | AF488 | ThermoFisher |
| Mouse |  | Goat PC | 1:250 | AF488 | ThermoFisher |

### Flow cytometry-based FRET experiments

HEK293T cells were seeded in a 12-well format one day prior to transfection. Two plasmids that encoded eCFP- or eYFP-tagged proteins of interest were used for transfection (Table 5). Cells were transfected with 1 µg of each plasmid using PEI as described above. Cells were harvested 24 h after transfection by detaching them with a pipet in 1 ml PBS. They were directly transferred to a 5 ml tube on ice. Cells were spun at 300 g for 5 min, supernatant was discarded, cell pellet was resuspended in 350 µl FACS buffer and cells were immediately measured for FRET signal using at a MACSquant VYB flow cytometer (Miltenyi Biotech).

### Split-Kusabira green experiments

HEK293T control and CD81KO (sorted for low CD81 expression) cells were transfected with plasmids encoding for fragments of p50 and p65 that were fused to parts of Kusabira green. 48h after transfection, cells were harvested as described above for flow cytometry-based FRET analysis and green fluorescence intensity was measured. Details are described in the CoralHue Fluo-chase Kit (MBL Int.corp.) manual.

### Immunofluorescence

Immunofluorescence experiments were performed in a 96-well format. Cells were fixed with 2% PFA for 10 min at RT before permeabilizing them with 80% acetone for 10 min at RT. Cells were then washed three times with PBS and blocked with 10% normal goat serum in PBS for 30 min at RT. After three washing steps, primary antibody (dilution 1:250) was applied for 1 h at RT, followed by three washing steps and secondary antibody incubation (dilution 1:250) for 1 h at RT. Then, nuclei were stained with DAPI (1:20,000, 10min, RT; Sigma Aldrich) or SiR-DNA dye (1 µM, 90 min, RT; Tebu-Bio). Three final washing steps were performed, and fluorescence was measured with a Cytation 3 plate imager (BioTek) or Incucyte plate imager (Sartorius).

### Luciferase assays

Luciferase assay experiments were performed in a 96-well format and measured in a 96-well white opaque plate using a Cytation 3 plate reader (BioTek) or a TriStar2 S microplate reader (Bertold Technologies). In case of Jc1_R2A Renilla luciferase reporter constructs, cells were electroporated with Jc1_R2A viral RNA and seeded in a 96-well plate. After incubation, cells were washed and lysed as described elsewhere (35). For NF-κB reporter activity, cells were transfected with several plasmids (Table 6). This included a NF-κB reporter plasmid (pNF-κB(3x)-FLuc), a *Gaussia* luciferase encoding plasmid as transfection control (pCMV-Gluc), and different plasmids encoding inducer proteins of NF-κB and IFN signaling cascades (p_human_IKKβ_ca, p(N)FLAG-CMV2 MAVS, pEF-Bos-RIG-I 1.211-flag). In some experiments, different chemical inducers of given pathways were used (phorbol 12-myristate 13-acetate (PMA; 10-100 ng/ml), TNFα (10 ng/ml), Lipopolysaccharide (LPS; 100 ng/ml), Ionomycin (0.25 µM), PolyIC (5 µg/ml, transfected)). Cells were additionally transfected with a control plasmid (pWPI_BLR) or a plasmid encoding CD81 (pWPI_hCD81-HAHA_BLR), and plasmids encoding eYFP or eYFP-core (pEYFP, pEYFP-HCV-core). 4 h after transfection, cells were treated with chemical inducers and luciferase activity was determined 24 h after transfection. For this, cell culture supernatant was transferred to a 96-well white opaque plate and mixed with coelenterazine (final conc. 5 µM) to determine transfection efficiency. Then, cells were washed with PBS and lysed with 60 µl FLuc lysis buffer (0.1 M KH_2_PO_4_/K_2_HPO_4_ (pH 7.8), 1% (v/v) Triton X-100, 1 mM DTT before use) for 10 min at RT. 40 µl lysate was transferred to a 96-well white opaque plate and mixed with 40 µl FLuc assay buffer (0.1 M KH_2_PO_4_/K_2_HPO_4_ (ph 7.8), 15 mM MgSO_4_, 5 mM ATP) and 40 µl FLuc substrate buffer (0.28 mg/ml D-Luciferin in FLuc assay buffer). Firefly luciferase signal was measured immediately.

### Microscopy

Live cell imaging was performed in a 96-well format using an Incucyte plate imager (Sartorius). Images were taken every 2-4 h in the respective channels. Imaging of fixed plates was performed using a Cytation 3 plate imager (BioTek) for cells stained for HCV core and with an Incucyte plate imager for cells stained for p65.

### Data Analyses

Design and alignment of DNA plasmids was done using SerialCloner v2.6.1 (SerialBasics) unless stated differently. Western blot membranes were analyzed usinge ImageStudio lite (LI-COR biosciences). Flow cytometry data was analyzed using Flowlogic v8.3 (Inivai Technologies). Microscopy images were analyzed using SoftWoRx 7.0 (Cytiva), Gen5 v3.10 (BioTek Instruments), IncuCyte GUI v2021A (Sartorius) according to instrument and subsequently handled with ImageJ. Statistical analysis and creation of graphs was done using GraphPad Prism 9 (GraphPad Spftware LLC) and Excel 2019 (Microsoft corp.). Arrangement of figures was done using CorelDraw X7 (Corel Corporation). The graphical abstract was created with BioRender.

## Results

### CD81 is downregulated in cells actively replicating HCV

In order to study alterations of the plasma membrane proteome upon active HCV replication, a flow cytometry-based surface expression screen was performed. In brief, Huh7.5 hepatoma cells were electroporated with viral genomic RNA, which encodes for a fluorescently labeled NS5A fusion protein (Jc1_NS5A-mtagBFP) to distinguish cells with active HCV replication (BFP+) from bystander cells (BFP-). Cells were then stained with an arrayed set of antibodies in a 96-well format, and the surface expression of 332 proteins was measured and compared between bystander and HCV-expressing cells, to calculate fold receptor modulation. Analysis revealed seven proteins whose levels were significantly lower upon HCV genome replication (Figure 1a). The two most down-regulated proteins, CD63 and the HCV entry receptor CD81 were independently confirmed upon electroporation of Huh7.5 hepatoma cells with viral genome RNA encoding for a fluorescently labeled NS5A fusion protein (Jc1_NS5A-eGFP). (Figure 1b) and chosen for further analysis. Notably, both are members of the family of tetraspanins. To get first insights on viral proteins involved in this phenotype, we transfected HEK293T cells to express single HCV proteins fused to eYFP and analyzed cell surface (Figure 1c) and total (Figure S1a) CD63 and CD81 levels by flow cytometry. Both, cell surface and less pronounced total tetraspanin levels were reduced upon expression of E2, NS5A and p7, indicating that the concerted action of several HCV proteins contributes to CD63 and CD81 downregulation.

**Figure 1:**
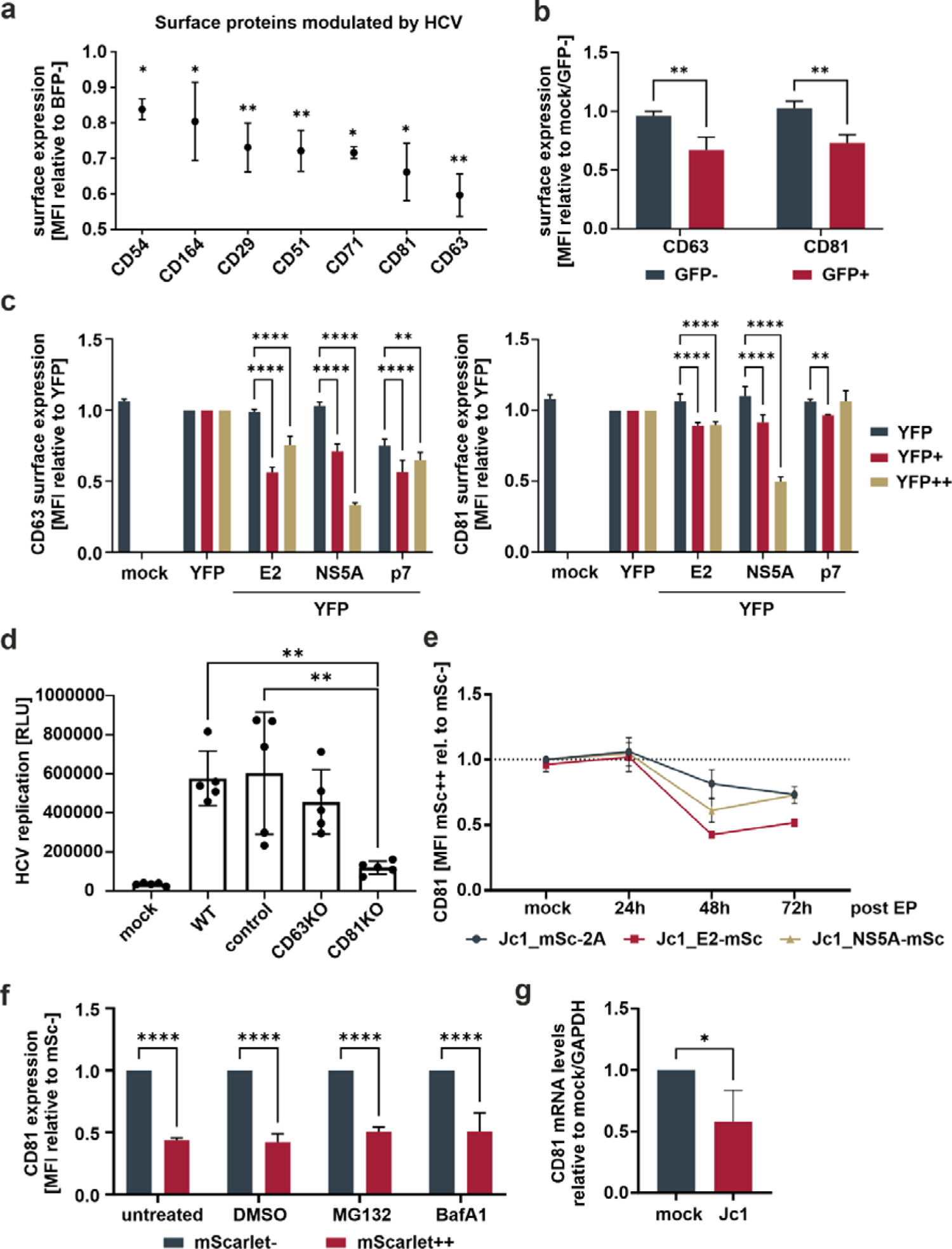
CD81 is actively downregulated in HCV expressing cells and interacts with various viral proteins. **(a)** Cell surface receptors significantly downregulated in HCV-expressing versus non-infected bystander cells. Huh7.5 cells were electroporated with Jc1_NS5A-mtagBFP HCV RNA and surface receptor expression screened with an arrayed panel of 332 PE-labeled antibodies 48h later by flow cytometry. Data from three independent screen. Shown are only receptors that were significantly modulated (p<0.05) in comparison to bystander (BFP-) cells. **(b)** Surface expression of tetraspanins CD63 and CD81 in HCV-expressing versus bystander cells. Huh7.5 cells were electroporated with Jc1_NS5A-GFP HCV RNA and cultivated for 72h. Cells were stained for tetraspanin surface expression and analyzed via flow cytometry. Depicted are data from 3 independent experiments. Significance was tested with a two-way ANOVA with Sidak’s multiple comparisons test. **(c)** Surface tetraspanin expression levels in cells transfected to express different viral proteins. HEK293T cells were transfected to express different YFP-tagged HCV proteins. 24h after transfection cells were harvested for flow cytometric analysis and stained for tetraspanins CD63 and CD81. Shown are transfection levels of bystander (YFP-) and cells expressing medium (YFP+) and high (YFP++) levels of viral proteins relative to YFP expressing cells. Depicted are data from 3 independent experiments. Significance was tested with a two-way ANOVA with Tukey’s multiple comparisons test. **(d)** Viral replication in Huh7.5 tetraspanin knock-out cells. Huh7.5 control and knock-out (KO) cells were electroporated with Jc1_R2A HCV RNA. 72h later cells were lysed and luciferase activity was measured as proxy for viral replication. Data from 5 independent experiments. Significance was tested with a one-way ANOVA with Tukey’s multiple comparisons test. **(e)** Relative total CD81 levels over time in cells expressing different viral genomes. Huh7.5 cells were electroporated with indicated viral RNAs. After different time points, cells were fixed, permebilized, stained for CD81 and measured by flow cytometry. Data from 2-4 independent experiments. **(f)** Total CD81 levels in cells treated with proteasomal or lysosomal inhibitors. Huh7.5 cells were electroporated with Jc1_NS5A-mScarlet and 48h later treated with either MG132 (1µM), Bafilomycin A1 (100nM) or DMSO (1%) as control. 24h after treatment cells were harvested, fixed, permeabilized, stained for CD81 and measured via flow cytometry. Data from 3 independent experiments. **(g)** CD81 mRNA levels in HCV and Mock electroporated cells. Huh7.5 cells were electroporated with Jc1 HCV RNA. 48h later, cellular mRNA was extracted and qRT-PCR was performed. Data from 4 independent experiments. Significance was tested with an unpaired t-test. All data show mean values ± SD if not mentioned otherwise. * p ≤ 0.05, ** p ≤ 0.01, *** p ≤ 0.001, **** p ≤ 0.0001.

To elucidate the functional role of tetraspanins for viral replication. For this, a HCV genomic RNA encoding a luciferase reporter was used (Jc1_R2A (31)). Hepatoma cells harboring gene knock-outs in CD63 and CD81 (Figure S1b) were electroporated to express Jc1_R2A, and luciferase activity was measured as a proxy for viral genome replication. As expected, given the role of C81 as HCV entry receptor, cells lacking CD81 showed a strongly decreased luciferase signal compared to controls or cells with CD63KO (Figure 1d).

As there was no apparent CD63 phenotype observable, we followed up on CD81 and analyzed the kinetic of total CD81 modulation in HCV-expressing cells. For this, we used an HCV reporter virus expressing the red fluorescent protein mScarlet, instead of the luciferase in front of the polyprotein (Jc1_mSc-2A), or Jc1-based constructs with internal E2 or NS5A mScarlet fusion proteins (Jc1_E2-mSc, and Jc1_NS5A-mSc, respectively). Monitoring mScarlet fluorescence allowed us to determine CD81 levels in HCV replicating and bystander cells over time (Figure1e, Figure S2a). While total CD81 levels were not altered 24h post EP, a decrease was observed at 48h which stayed similar at 72h. Since not only cell surface, but total CD81 levels were reduced, we investigated if CD81 is actively degraded by HCV. We electroporated Huh7.5 cells with Jc1_NS5A-mScarlet, treated them with a proteasomal (MG132) or a lysosomal inhibitor (Bafilomycin A1), stained for total CD81 levels and analyzed CD81 modulation by flow cytometry. Treatment with none of the inhibitors rescued CD81 levels indicating that CD81 is not degraded by the proteasome or the lysosome in HCV-expressing cells (Figure 1f). Instead, RT-qPCR revealed that the levels of CD81 mRNA were reduced by approximately half 48 h after electroporation of Huh7.5 cells (Figure1g). Of note, we here used non-modified HCV JC1, demonstrating that CD81 is also modulated by non-tagged viral genomes. Taken together, HCV downregulates CD81 at the mRNA level in Huh7.5 cells involving several viral proteins, in particular E2, NS5A and p7.

### CD81-deficient hepatoma cells support HCV-expression and cell growth

CD81 is transcriptionally silenced and downregulated upon the onset of viral genome replication (Figure 1e and g). This suggests, that downregulation of CD81 plays additional roles in HCV biology beyond serving as entry receptor. A common strategy of viruses is to prevent superinfection by downmodulating its entry receptors and this has been described for HCV and CD81 (36). In order to analyze if reduction of CD81 by HCV confers additional benefits in the viral life cycle, we took advantage of HCV reporter genomes producing virions that are either severely (Jc1_NS5A-mScarlet) or completely (Jc1_E2-mScarlet) compromised in their ability to de novo infect cells. Infectious Jc1_mScarlet-2A was used as a positive control and live cell imaging over a period of seven days revealed, as expected, that this virus was not able to spread in CD81-negative Huh7.5 cells (Figure 2a, left). In contrast, Jc1_NS5A-mScarlet and Jc1_E2-mScarlet failed to induce spreading infection in both, control as well as CD81KO Huh7.5, confirming their loss of infectivity (Figure 2a, middle and right panel). However, of note, we consistently observed higher numbers of HCV-expressing cells in the CD81KO cells in comparison to the CD81-positive Huh7.5 controls (Fig. 2a, middle and right panel). Moreover, this phenotype was recapitulated in CD81-negative Huh7 Lunet cells that were engineered to express hCD81 (Fig. 2b). Similar to the conditions in which CD81 was knocked-out in Huh7.5, the number of HCV Jc1_E2-mScarlet replicating cells was high in CD81-negative parental Lunet cells and reduced upon reconstitution of hCD81. In addition, cellular proliferation and growth was reduced in Lunet cells upon expression of hCD81 (Fig.2b). In conclusion, CD81 negatively regulates HCV-replication and cellular growth.

**Figure 2:**
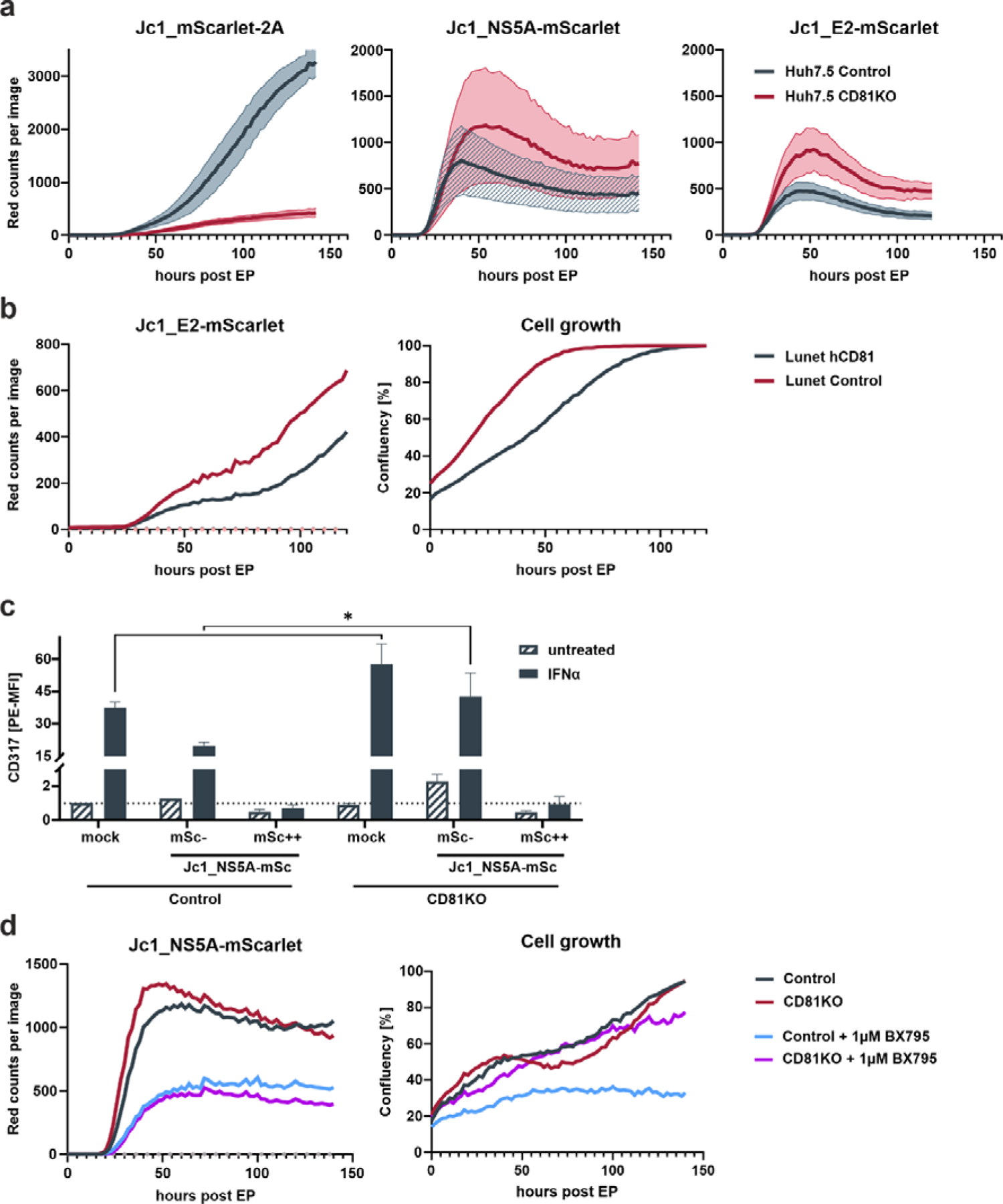
Loss of CD81 promotes growth and survival of HCV-expressing cells. **(a)** Replication kinetics of different fluorescently-labeled HCV genomes in Huh7.5 control and CD81KO cells. Huh7.5 control and CD81KO cells were electroporated with indicated viral genomes and fluorescence was measured over time (every 2h). Representative data from three independent experiments where lines represent mean values of images (4 images per well, three wells per condition) and their SD in semi-transparent area. **(b)** Cell growth and number of HCV expressing cells is reduced in CD81-expressing Huh7 cells. Huh7-Lunet and Huh7-Lunet-CD81 cells were electroporated with Jc1_E2-mScarlet. HCV expressing cells (red fluorescence count) and confluency was monitored over time. Shown is one representative experiment. **(c)** ISG counteraction by HCV is independent of CD81. Huh7.5 control and CD81KO cells were electroporated with Jc1_NS5A-mScarlet. 48h after EP cells were treated with IFNα (10ng/ml) for additional 24h, then harvested for flow cytometry and stained for surface expression of tetherin (CD317). Shown are tetherin surface experssion levels relative to untreated mock electroporated Huh7.5 control cells. mSc-represent bystander cells while mSc++ represent cells highly expressing NS5A-mScarlet. See also Figure S3A. Data from 2 independent experiments with tripilicate infection. Significance was tested with a two-way ANOVA with Sidak’s multiple comparisons test), **(d)** Loss of CD81 compensates for pro-survival TBK1 signalling in HCV-expresing cells. Huh7.5 control and CD81KO cells were electroporated with Jc1_NS5A-mScarlet and treated with BX795 (1µM). HCV expressing cells (red fluorescence count) and confluency was monitored over time. Shown is one representative of 2 independent biological replicates..

### CD81 reduces pro-survival signaling in HCV-expressing hepatoma cells

The network of HCV protein interactions was generally not altered in CD81-depleted cells (Figure S2b), indicating that other factors are responsible for the inhibitory effect of CD81 on the growth of HCV-expressing cells. Interferon (IFN) treatment triggers an antiviral state that renders cells largely resistant to HCV replication. Nevertheless, cells that actively replicate HCV genomes can overcome early IFN-mediated antiviral immune response by expression of viral proteins that counteract the expression of IFN-regulated genes (ISGs) (37). We hypothesized that CD81 might possibly alter IFN-signaling and thereby restrict intracellular HCV replication. Therefore, Huh7.5 control and CD81 KO cells were electroporated with Jc1_NS5A-mScarlet, stimulated with IFNα and stained for the interferon-stimulated gene (ISG) tetherin (CD317), to analyze if Huh7.5 are responsive to IFNα despite the lack of PRRs and if HCV counteracts this response in our system (Fig. 2c). Mock electroporated cells showed a clear induction of tetherin expression 24 h after IFNα stimulation. A similar effect was observed for Jc1_NS5A-mScarlet electroporated cells that were mScarlet-negative and did hence not express viral proteins (bystander cells, gated as mSc-, 2c, **Error! Reference source not found.**). In contrast, cells that actively replicate HCV as evident by NS5A-mScarlet expression, had tetherin levels comparable to cells that were not treated with IFNα, demonstrating that the IFN-mediated antiviral immune response is indeed suppressed in HCV-expressing cells (2c, **Error! Reference source not found.**). In addition, CD81 did not impair the ability of HCV to counteract the IFN-response, as tetherin induction was suppressed in control cells to a similar extent as in CD81KO Huh7.5. A remarkable difference was baseline induction of tetherin upon IFNα treatment. CD81KO cells that were mock electroporated or the mScarlet-negative bystander cells, both expressed higher tetherin levels as control cells, indicating that the absence of CD81 could sensitize cells for ISG induction (Figure 2c, Figure S3).

The interferon signaling cascade involves activation of TBK1, which has a central role in innate immunity and ISG induction. In addition, TBK1 is involved in pro-survival and anti-apoptotic signaling and can activate NF-κB as downstream target (38). As HCV blunts ISG induction, pro-survival signaling of TBK1 could be beneficial for viral replication and growth of HCV-expressing cells. To test for this, we electroporated Huh7.5 control and CD81KO cells with Jc1_NS5A-mScarlet and treated them with a TBK1 inhibitor (BX795). Of note, inhibition of TBK1 led to decreased viral replication in both, Huh7.5 control and CD81KO cells (Figure 2d, left). However, only Huh7.5 CD81KO cells proliferated after being electroporated with HCV RNA and treated with BX795 (Figure 2d, right, purple line as compared to blue line). This suggests that the absence of CD81 can compensate for TBK1-mediated inhibition of proliferation of Huh7.5 hepatoma cells.

### CD81 is a negative regulator of NF-κB

We hypothesize that TBK1 induces pro-survival signaling via NF-κB that is suppressed by CD81. Furthermore, NF-κB is activated via HCV Core inducing proliferation of hepatoma cells and possibly tumor formation (39–44). To study effects of Core and CD81 on NF-κB we first used HEK293T transfected to express a NF-κB luciferase reporter plasmid together with a control or CD81 plasmid and induced signaling by co-transfection of HCV YFP-core plasmid with or without TNFα treatment. Indeed, HCV Core and TNFα induced NF-κB reporter activity ∼20-fold, while both together led to a ∼60-fold induction (Figure 3a). Of note, transfecting cells to express CD81 reduced NF-κB reporter activity induced by HCV Core and TNFα nearly to background levels.

**Figure 3:**
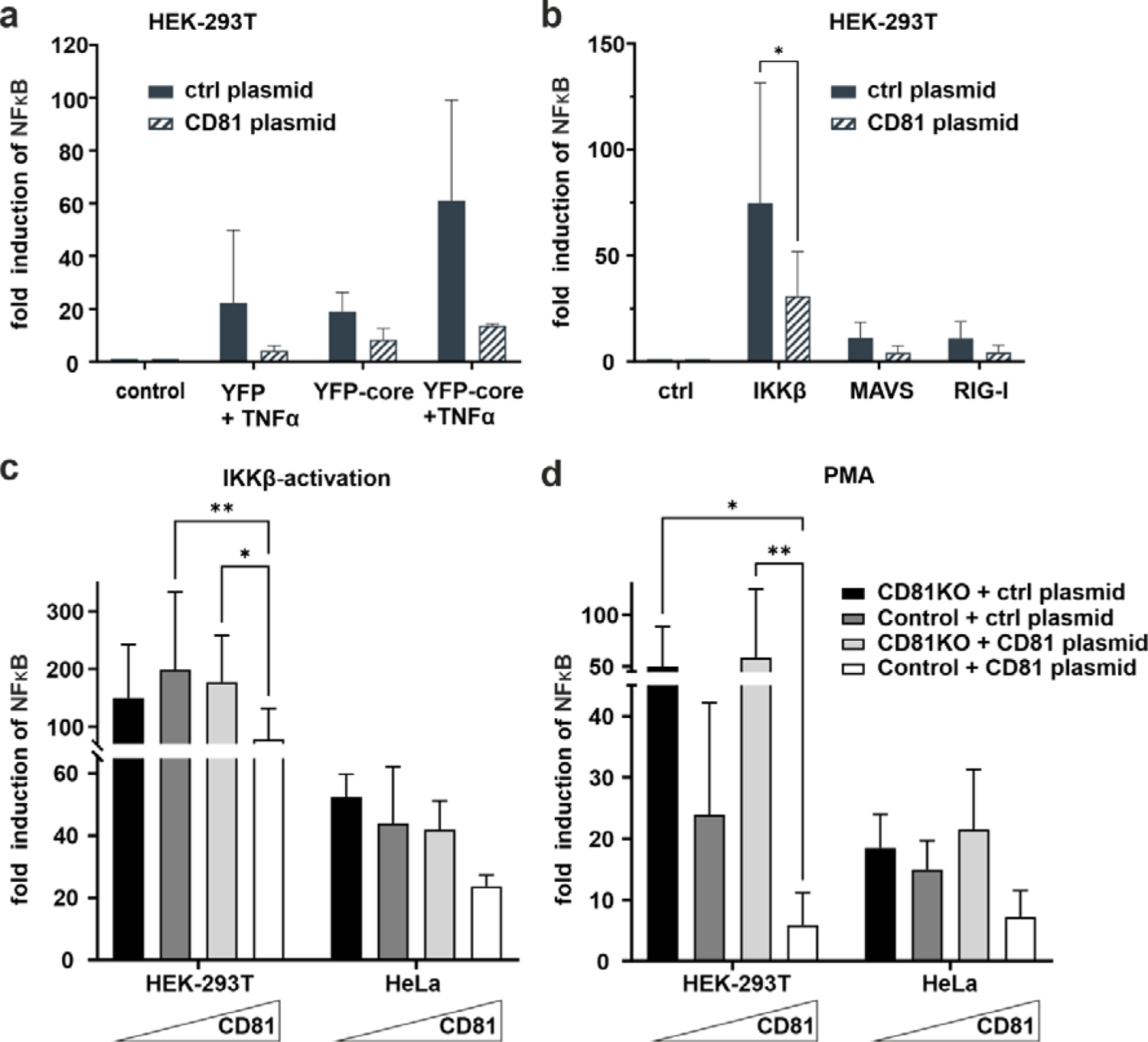
CD81 suppresses IKKβ and PMA-mediated NFκB activation. **(a)** CD81 suppresses NFκB activation by TNFa and HCV core. HEK293T cells were transfected to express a NFκB luciferase reporter and with a plasmid encoding CD81 or a control plasmid (control vector). Additionally, cells were transfected with HCV core (YFP-core) or a control plasmid (YFP alone). 4h after transfection, cells were treated with TNFa (10ng/ml). 24h after transfection cells were lysed and luciferase activity was measured. Data from 2 independent experiments. (**b**) CD81 reduces NFκB activation by IKKβ, MAVS and RIG-I. HEK293T cells were transfected to express a NFκB luciferase reporter and plasmids encoding either IKKβ, MAVS or RIG-I. Additionally, a plasmid encoding CD81 (or empty control vector) was transfected. Data from 4 independent experiments. Significance was tested with a two-way ANOVA with Sidak’s multiple comparisons test. **(c and d)** CD81 reduces NFκB activation induced by IKKβ and PMA. HEK293T and HeLa control or CD81KO cells were transfected to express a NFκB luciferase reporter. Additionally, a plasmid encoding CD81 (or empty control vector) was transfected. Either an IKKβ encoding plasmid (**c**) was transfected together with the other plasmids, or **(d)** cells were treated with PMA (10ng/ml) 4h after transfection. 24h after transfection cells were lysed and luciferase activity was measured. Data from 4 independent experiments. Significance was tested with a two-way ANOVA with Tukey’s multiple comparisons test. All data points show mean values ± SD. * p ≤ 0.05, ** p ≤ 0.01.

We next sought to more closely investigate effects of CD81 on NF-κB signaling induced by either IKKβ, MAVS or RIG-I. Activity of the NF-κB reporter was induced by all of them, with IKKβ being the most potent inducer and again upon expression of CD81 in this system, a clear reduction of NF-κB activity could be detected (Figure 3b). Subsequently, we decided to analyze cell type dependency by using HEK293T as well as HeLa cells, varied CD81 levels by employing cells with and without CD81KO that were additionally transfected to express CD81 or a control plasmid and used Phorbol 12-myristate 13-acetate (PMA) as an exogenous NF-κB trigger in addition to IKKβ (Figure 3c and d). In general, irrespective of the cell line or inducer used (IKKβ or PMA), increasing amounts of CD81 reduced NF-κB reporter activity, albeit the effect was only significant in HEK293T cells (Figure 3c and d).

We next thought to corroborate our results by more closely investigating CD81KO HEK293T and HeLa cells and induce NF-κB exogenously via PMA, instead of transfecting IKKβ (Figure 4). PMA mimics the second messenger lipid diacylglycerol (DAG) which activates protein kinase C (PKC). Some PKC isoforms require the second messenger Ca^2+^, to become fully activated (45). Hence, cells transfected with the NF-κB reporter were additionally treated with Ionomycin to increase cytosolic Ca^2+^ levels. Ionomycin treatment did not increase NF-κB activity and did not induce NF-κB activity when administered alone (Figure 4a and b). However, in line with our previous results, NF-κB reporter activity in HEK293T (Figure 4a) and HeLa cells (Figure 4b) was strongly suppressed by CD81 and we also observed suppression of PMA-induced NF-κB reporter activity by endogenous CD81 (Fig. 4c). Using a more physiological relevant readout in non-transfected HEK293T control and CD81KO cells, we also found that levels of TNFα mRNA were increased at different time points post PMA stimulation, when cells were depleted for CD81 (Figure 4d).

**Figure 4:**
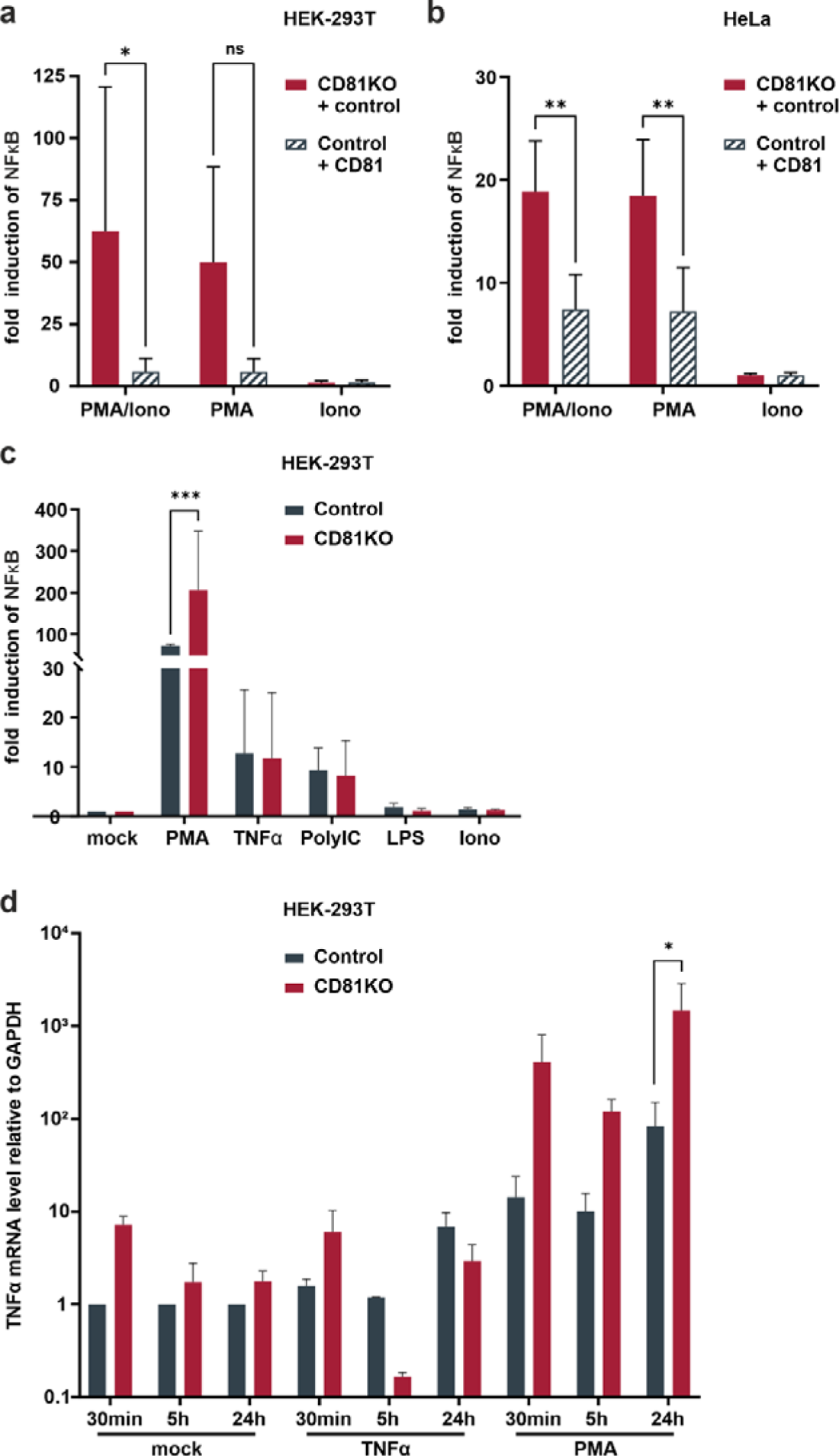
Suppression of NFκB transcriptional activity by CD81 following PMA-stimulation. CD81 suppresses PMA-mediated NFκB activation in different cell lines. **(a)** HEK293T or **(b)** HeLa control and CD81KO cells were transfected to express a NFκB luciferase reporter. Additionally, cells were transfected with a plasmid encoding CD81 or a control plasmid (empty control vector). 4h after transfection cells were treated with PMA (10ng/ml), Ionomycin (0.25µM), or both. 24h after transfection cells were lysed and luciferase activiy was measured. Data from 4 independent experiments. Significance was tested with a two-way ANOVA with Tukey’s multiple comparisons test. **(c)** HEK293T control and CD81KO cells were transfected to express a NFκB luciferase reporter. 4h after transfection cells were treated with PMA (10ng/ml), TNFa (10ng/ml), LPS (100ng/ml), Ionomycin (0.25µM), or PolyIC (5µg/ml, transfected with plasmids). 24h after transfection cells were lysed and luciferase activity was measured. Data from 3 independent experiments. Significance was tested with a two-way ANOVA with Sidak’s multiple comparisons test. **(d)** NFκB transcriptional activity is increased in CD81KO cells. HEK293T control and CD81KO cells were treated with PMA (10ng/ml), TNFa (10ng/ml) or left untreated (Mock) for indicated time periods. Then, cellular RNA was extracted and TNFa mRNA was quantified via qRT-PCR. Data from 4 independent experiments as mean values ± SEM. All other data mean values ± SD. * p ≤ 0.05, ** p ≤ 0.01m *** p ≤ 0.001.

We next explored different potential mechanisms of increased NF-κB signaling and found that the interaction of the NF-κB signaling components p50 and p65 was increased in CD81 KO vs control HEK293T cells (Figure 5a). Furthermore, moving to hepatoma cells, we found that basal levels of p65 and phosphorylated p65 were strongly elevated, even though independently of TNFα (Figure 5b). To unambiguously directly analyze endogenous NF-κB signaling in Huh7.5 cells, nuclear translocation of p65, a subunit of the NF-κB transcription factor family, was followed upon treatment with PMA or TNFα (Figure 5c). PMA treatment induced p65 translocation after approximately 30 min which was strongly increased in CD81KO hepatoma cells (Figure 5c). Quantification of the nuclear intensity of the p65 staining revealed that while it reached a plateau in control cells at 30 min after treatment, it further increased in CD81KO Huh7.5 from 60 min post treatment on (Figure 5d). A similar trend was observed for TNFα-treated samples (Figure 5c and d). In conclusion, the cumulated results employing various NF-κB inducers, cell lines, overexpression and knockout conditions as well as different NF-κB readouts based on reporters and endogenous signals suggests a suppression of NF-κB activation by CD81.

**Figure 5:**
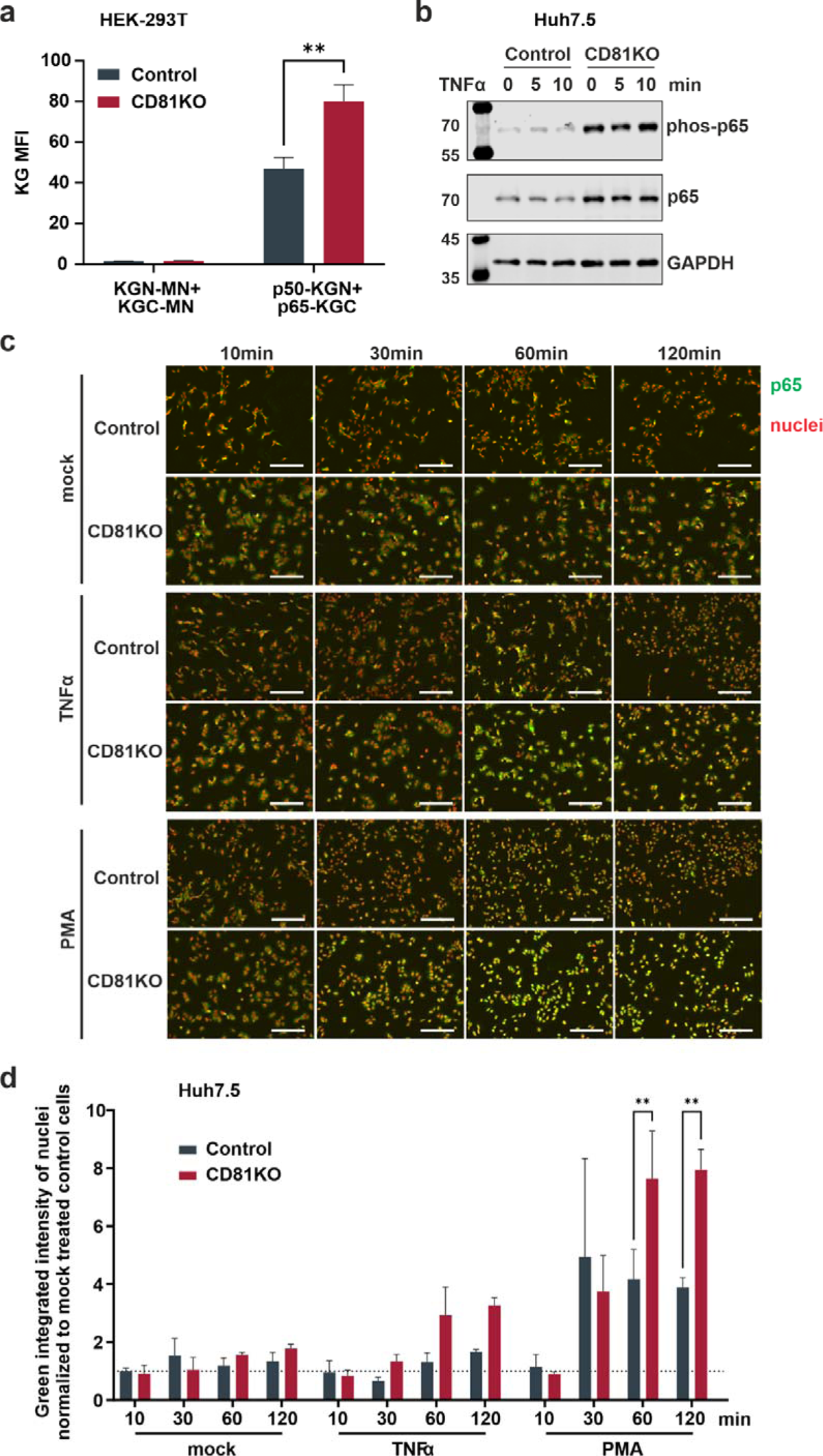
CD81 suppresses p65 nuclear translocation. **(a)** Interaction of p50 and p65. HEK293T CD81 KO and control cells were transfected to express p50 or p65 fused to fragments of Kusabira green (KGN and KGC, respectively). 48 h after transfection cells were harvested for flow cytometry. Close proximity allows reconstitution of full Kusabira green and fluorescence, indicating interaction. Depicted are data from 3 independent experiments. Significance was tested with a two-way ANOVA with Sidak’s multiple comparisons test. ns = not significant, * p ≤ 0.05, ** p ≤ 0.01. **(b)** Protein amount and phosphorylation status of p65 in Huh7.5 cells. Huh7.5 control and CD81KO cells were treated with TNFa (10ng/ml) for indicated time points. Then, cells were lysed and subjected to SDS-PAGE and western blot. Shown is one representative experiment. **(c)** Representative images of p65 nuclear translocation after stimulation. Huh7.5 control and CD81KO cells were mock treated, with TNFα (10ng/ml) or PMA (10ng/ml) for the indicated time periods. Then, cells were fixed, permeabilized and stained for p65 (green) and nuclear DNA (red). Shown are representative images from 3 independent experiments. **(d)** Quantification of p65 nuclear translocation after stimulation. Shown is the green integrated intensity (representing p65) in areas that overlap with red fluorescence (representing nuclear DNA) relative to unstimulated cells. Data from 3 independent experiments as mean values ± SEM. Significance was tested with a two-way ANOVA with Sidak’s multiple comparisons test.

## Discussion

Here, we identify CD81 as repressor of pro-survival signaling in hepatoma cells by interfering with NF-κB activation. Apart from elucidating a novel thus far unknown cellular function of the tetraspanin CD81, this is an intriguing mechanism of how HCV manipulates infected cells to achieve persistent and chronic infection. Lower levels of CD81 in cells infected with HCV or cells that stably express viral proteins have been observed before and were suggested to play a role in superinfection exclusion (36,46–48). Indeed, downregulation of the main entry receptor is a common strategy of viruses to prevent superinfection (49,50). However clearly, superinfection exclusion is not a denominator of the effects observed here, as CD81 is downregulated post entry and the positive phenotype of CD81-depletion on proliferation of HCV-expressing cells was observed using tagged HCV-reporter genomes that produce non-infectious virions. Furthermore, the repression of NF-κB via CD81 is completely independent of HCV-expression. It is thus remarkable, that by lowering levels of CD81, HCV independently mediates superinfection exclusion as well as enhancement of pro-survival intracellular signaling – two mechanisms that are highly likely to promote chronic and persistent infection. Indeed, and in line with our hypothesis, it was suggested previously that in the context of persistent HCV infection there is a selection towards maintenance of CD81-low cells, that are more resistant to cell death and apoptosis (36). We corroborate this hypothesis and extend it to mechanistic functions of CD81 in suppressing NF-κB signaling. Apart from that, it is noteworthy, that apparently from 332 surface receptors included in our screen, only seven were significantly lower in HCV-expressing cells (Figure 1a), which is in contrast to other viruses i.e. HCMV and HIV (27,51,52), that heavily dysregulate the plasma membrane of infected cells for evasion of antiviral immune responses. This indicates that HCV, for efficient persistent and chronic infection adopts a “stealth” mode in infected cells, instead of blunting adaptive and humoral cellular immune responses by cell surface receptors dysregulation.

While it is clear that HCV is generally sensitive to IFN and cannot overcome the antiviral state in the context of *de novo* infection, it highly efficiently blunts interferon signaling and thereby innate antiviral immune responses in actively replicating cells (Figure 2c) (37,53–55). In such a scenario, NF-κB activation is pro-survival in the absence of innate immune activation (56), which could explain the inhibitory phenotype of HCV replicating cells when blocking TBK1 (Figure 2d).

A variety of interesting and important questions remain currently unanswered. Concerning the mechanism of CD81 downregulation by HCV, our data is in line with previous work reporting transcriptional silencing in cells stably expressing HCV NS4B (46). In our study, CD81 levels were mainly affected by NS5A and to a lesser extent by E2 and p7 (Figure 1c and S1a). Unfortunately, transient expression of NS4B was weak in our experiments, which is why we had not analyzed this viral protein. However, it is conceivable and highly likely that HCV evolved various mechanisms to downregulate CD81 as one of its main entry receptors. Similarly, for instance HIV, uses Nef, Vpu and Env to downregulate the primary receptor CD4 (57–61). Altogether, the cumulated data of us and our colleagues (46–48) establishes an important role of CD81 in HCV biology beyond serving as main entry receptor, that warrants further investigation.

Similarly, it will be highly important to decipher how exactly CD81 suppresses NF-κB signaling For instance it is known that there is a mechanistic interplay of the integrated stress response and NF-κB (62). In this context, Fink et al. revealed that the activity of the ISR component IRE1α is important for HCV replication as it regulates cell survival, presumably by degrading the pro-apoptotic miR-125a (63). They further showed that knock-out of another ISR factor, that is XBP1, with simultaneous activation of IRE1α by NS4B renders cells resistant to the intrinsic pathway of apoptosis. Together with the study of Tardif et al., this connects higher ISR activity to pro-survival NF-κB signaling (63,64).

CD81-mediated suppression of NF-κB was most prominent in PMA-stimulated cells indicating that CD81 represses mainly this signaling pathway. The downstream target of PMA are PKCs that are activated by diacylglycerol (DAG) which is mimicked by PMA. This stimulus is combined with ionomycin, as some PKC isoforms are only activated in the presence of Ca2+, released by Ionomycin (45). However, Ionomycin was dispensable for NF-κB activation in our experimental settings indicating that CD81 suppresses the NF-κB signaling cascade by interfering with a subfamily of novel Ca^2+^ independent PKCs (45). A hypothesis that requires further experimentation, but is supported by the fact that the serotonin receptor of the 5-HT2 family signals through DAG and PKC and interacts with CD81 (65,66). In the context of HCV, Core induced NF-κB with TNFα (67). We confirm this mechanistic interplay and further demonstrate that CD81 efficiently suppresses this activation (Figure 3a). Given the importance of NF-κB for HCV replication and persistence, as well as its role in tumor development and hepatocellular carcinoma (39–44), it is tempting to speculate that HCV-mediated downregulation of CD81 is a thus far unprecedented mediator of viral tumorigenesis (14,68–71). Indeed, low levels of CD81 correlate with HCC metastasis and tumor proliferation (72,73) and expression of CD81 suppresses hepatocellular carcinoma development (74). Importantly, the role of CD81 in tumor development seems multifaceted with suppressive as well as protooncogenic functions (75) and could be dependent on the tumor type as well as co-factors, for instance HCV that interferes with innate immune signaling in infected hepatocytes.

Our study has certain limitations that need to be addressed in future work. First of all, HCV exerts a large genotype and subtype dependent heterogeneity and we only addressed CD81-modulation by the Jc1 viral genome, which is derived from an acute fulminant hepatitis (76). Hence, it will be exciting to address if CD81-modulation is a conserved feature of highly variable HCV genomes. Furthermore, even though we have witnessed CD81-dependent suppressive effects on NF-κB in different settings, including overexpression and KO as well as endogenous p65 translocation in different cell types (HeLa, 293T and Huh7.5) we have not assessed primary hepatocytes thus far, to verify this phenotype also in a non-tumorigenic setting. Finally, the mechanistic details, as discussed above, remain largely elusive and need to be carefully investigated. Nevertheless, altogether, we here identify unprecedented roles of CD81 in the context of HCV infection biology, pathogenesis and beyond.

## Consent for publication

All authors gave their consent to publish. All authors read and approved the final manuscript.

## Availability of data and material

All data generated and analyzed during this study are included in this published manuscript.

## Authors’ contributions

MB performed most of the experiments supported by ME, DH and MR and JN. MB and MS planned the experiments and analyzed the data. MB and MS wrote the manuscript draft. MS supervised the overall study and provided resources. All authors contributed to editing and developed the manuscript to its final form.

## Acknowledgements

We thank Gisa Gerold, Thomas Pietschmann and Daniel Sauter for the kind contribution of cell lines, reagents and plasmids and Daniel Sauter for critical reading of the manuscript

## Supplemental Figures

**Figure S1:**
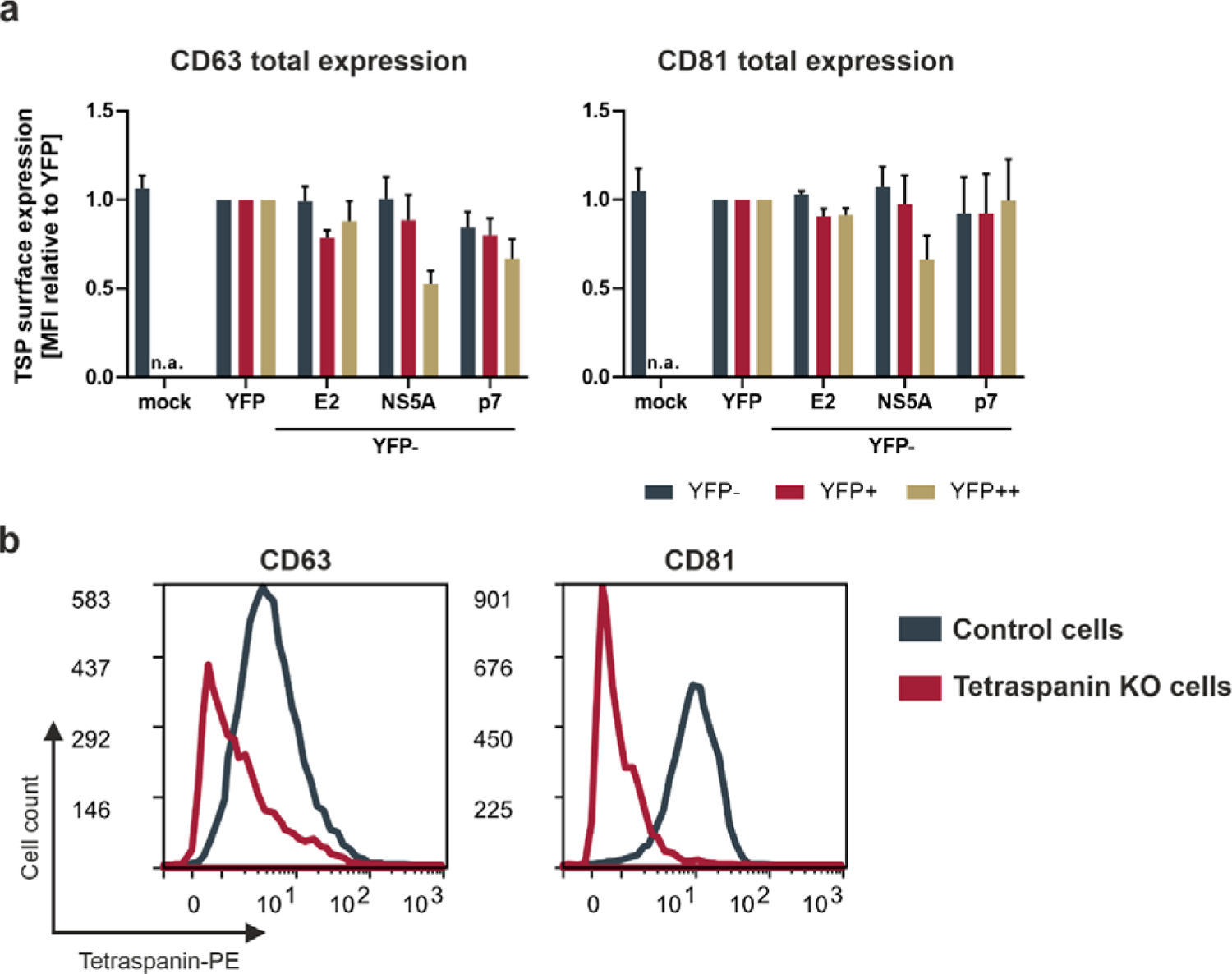
Downregulation of tetraspanins by HCV viral proteins and tetraspanin knock-out verification. **(a)** Total tetraspanin expression levels in cells transfected to express different viral proteins. HEK293T cells were transfected with plasmids encoding different YFP-tagged HCV proteins. 24h after transfection cells were harvested for flow cytometric analysis and stained for tetraspanins CD63 and CD81. Shown are transfection levels of bystander (YFP-) and cells expressing medium (YFP+) or high (YFP++) levels of viral proteins relative to YFP expressing cells. Data from 3 independent experiments. **(b)** Histograms of a flow cytometric analysis of tetraspanin knock-out cells for CD63 and CD81 cell surface expression. Huh7.5 cells were transduced with lentiviral particles that stably introduce a cassette encoding guide RNA and Cas9, together with a puromycin resistance into the host cell genome. Cells were selected for at least 2 weeks with puromycin (1µg/ml).

**Figure S2:**
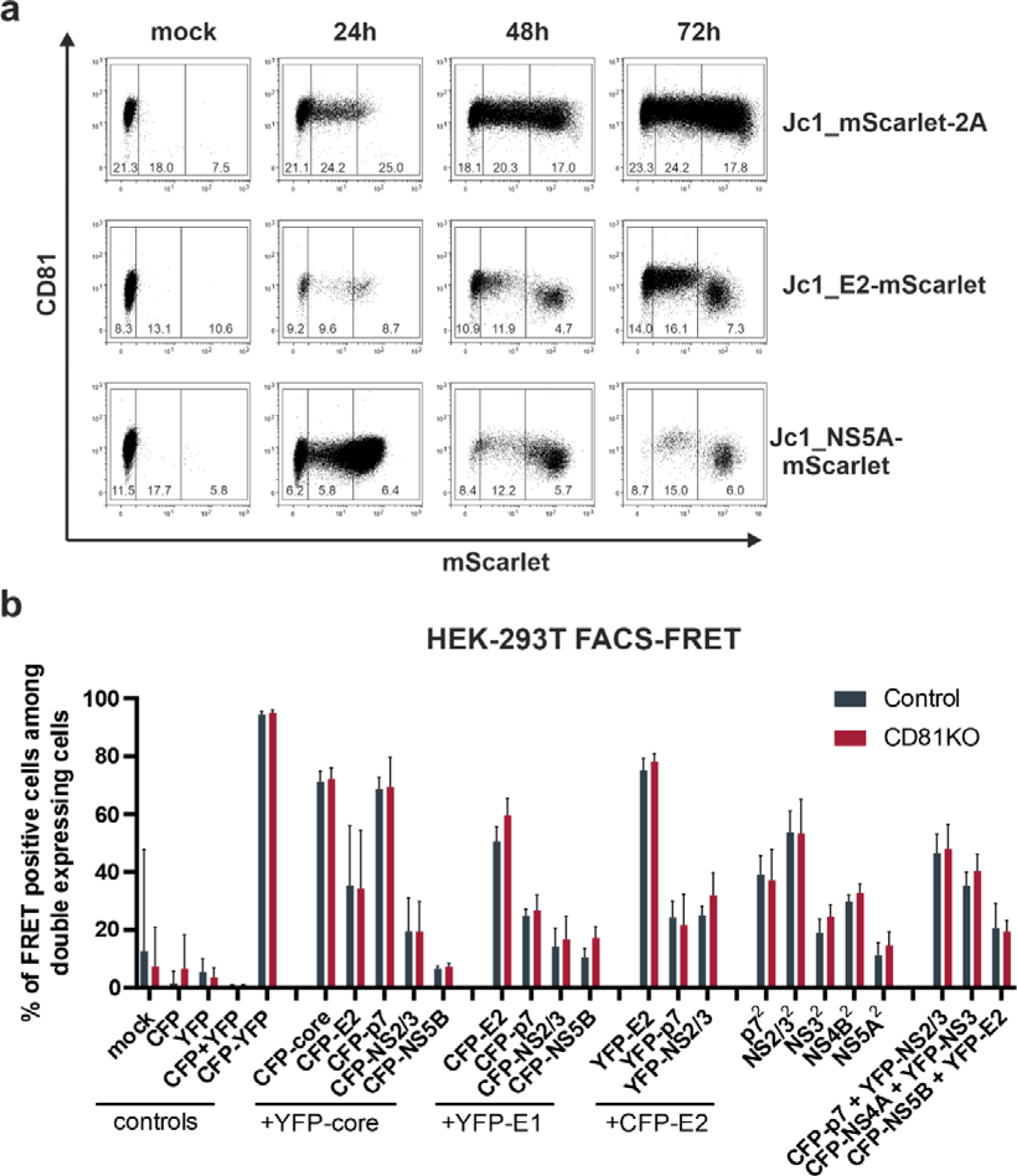
CD81 downmodulation and protein interactions of HCV proteins. **(a)** Representative experiment showing CD81 downregulation by different viral genomes. Huh7.5 cells were electroporated with indicated viral genome RNAs and harvested for flow cytometric analysis at indicated time points. Cells were permeabilized and stained for CD81. Gates show bystander (mSc-; left gate), medium (mSc+; middle gate) and highly (mSc++; right gate) HCV expressing cells. Numbers within gates show mean flourescent intensity (MFI) of CD81 expression. **(b)** Interaction network of viral proteins with each other. HEK293T control and CD81KO cells were transfected to express a pair of eCFP- and eYFP-tagged viral proteins. 24h after transfection cells were harvested and FRET signals were measured via flow cytometry. Only combinations that showed interaction in a previous study were tested *(77)*. Data from 4-8 independent experiments, mean values ± SD.

**Figure S3:**
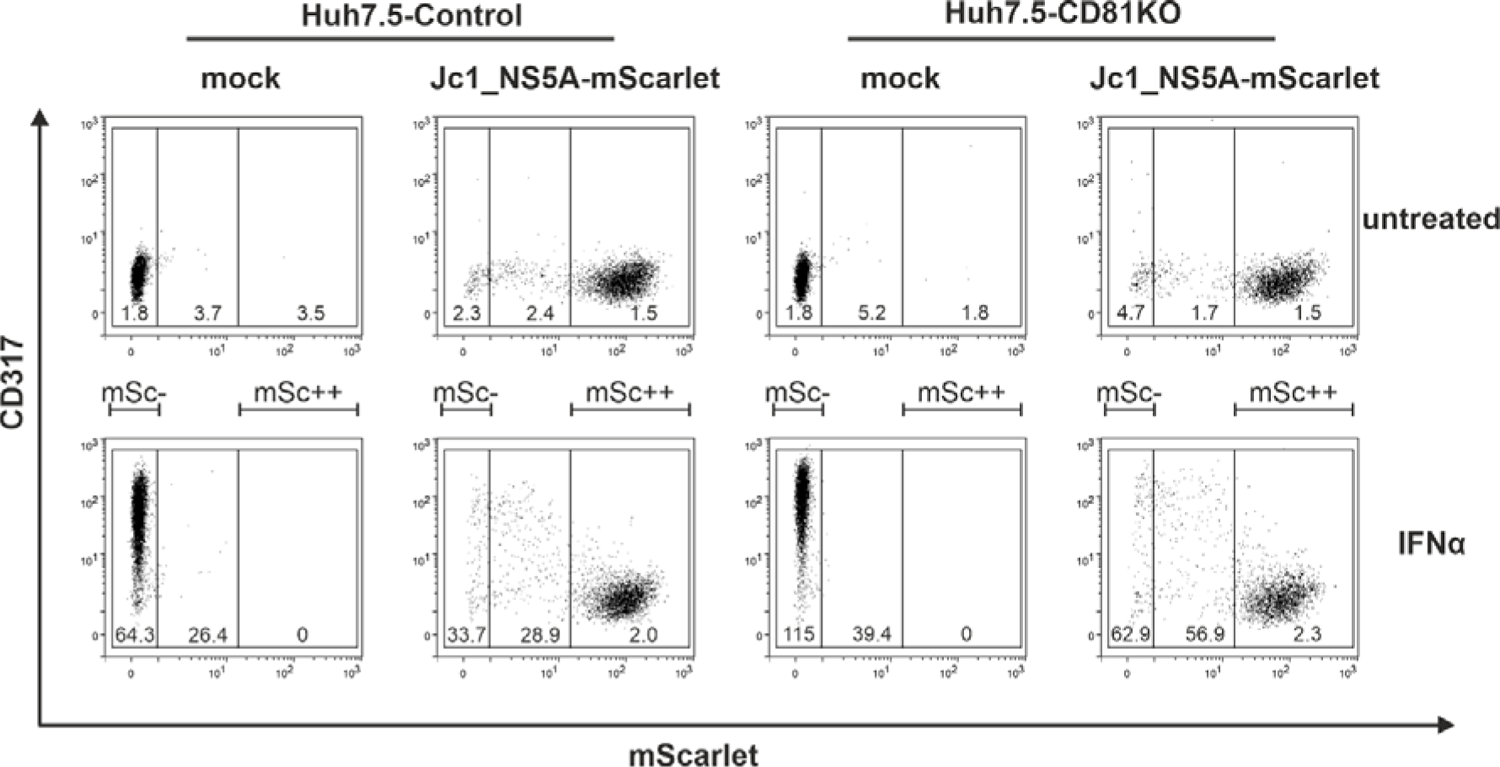
CD81 does not impair the ability HCV to counteract the interferon response. Representative experiment of HCV expressing cells treated with IFNa, see also Figure 2c. Huh7.5 control and CD81KO cells were electroporated with Jc1_NS5A-mScarlet. 48h after EP cells were treated with IFNα (10ng/ml) for additional 24h, then harvested for flow cytometry and stained for surface expression of the ISG tetherin (CD317) as proxy for the interferon response. Gates show bystander cells (mSc-; left gate), and cells that express high (mSc++; right gate) levels of NS5A-mScarlet. Numbers within gates represent mean fluorescence intensity (MFI) of tetherin (CD317) expression.

